# A Three-Dimensional Bioprinted Skin Melanoma Model for High-Throughput Drug Screening

**DOI:** 10.64898/2026.11.06.764888

**Authors:** Paige Derr, Kristy Derr-Mierzwa, Shayne Frebert, Min Shen, Ty Voss, Marc Ferrer

## Abstract

Melanoma is the deadliest form of skin cancer, and despite major advances in immunotherapy and targeted therapies, most patients with advanced disease eventually develop treatment resistance. The lack of physiologically relevant, scalable preclinical models that recapitulate the tumor microenvironment (TME) remains a major barrier to identifying more effective therapies. Here, we present a novel high-throughput screening platform based on a bioprinted, full-thickness human melanoma skin model that combines the biological complexity of native skin with the scalability required for drug discovery. The 3D tissue construct comprises a dermal region containing primary human dermal fibroblasts, melanoma tumor-like cell clusters localized at the dermal-epidermal interface, and an overlying differentiated epidermis composed of primary human keratinocytes cultured at the air-liquid interface, thereby recapitulating key structural and cellular features of the melanoma TME. To enable multiplexed, high-content analysis, melanoma cells and fibroblasts were engineered to express red and green fluorescent proteins, respectively, allowing automated confocal imaging to simultaneously quantify tumor growth and stromal viability. This dual-readout strategy provides an intrinsic counterscreen to distinguish melanoma-selective compounds from nonspecific cytotoxic agents. The assay was implemented in a 96-well bioprinted format compatible with high-throughput screening of compound libraries across multiple concentrations. Using this platform, we identified compounds that selectively reduced melanoma cell viability while sparing surrounding stromal cells, demonstrating the value of incorporating the TME into primary drug screens. By integrating bioprinting, human tissue engineering, and automated high-content imaging into a scalable workflow, this platform provides a robust and clinically relevant approach for melanoma therapeutic discovery and has the potential to improve the translation of preclinical findings into effective treatments for advanced melanoma.

## Introduction

Melanoma is one of the most aggressive and rapidly increasing malignancies worldwide, and although it accounts for only a small fraction of all skin cancers, melanoma is responsible for most skin cancer-related deaths [1,2]. Incidence rates have risen rapidly over recent decades, increasing faster than those of almost any other cancer type, particularly among populations with fair skin and high ultraviolet radiation exposure [3]. Melanoma treatment is highly personalized, depending on the cancer stage and tumor genetics. Primary options include surgical removal, immunotherapy, targeted therapy, and radiation. Early-stage melanoma can often be cured with surgery alone, while advanced or metastatic cases rely heavily on systemic medications. While recent advances in immunotherapy and targeted therapy have improved survival for a subset of patients, five-year survival rates for stage IV disease remain below 30%, and many patients develop resistance to currently available agents [4,5]. Developing better treatments for advanced and metastatic melanoma is a critical unmet medical need.

In cancer, the tumor microenvironment (TME) plays a major role in growth, expansion, and resistance to treatment. In melanoma, the TME induces resistance to some of the treatments, including immunotherapies like checkpoint inhibitors (e.g., anti-PD-1, anti-CTLA-4). The development of new therapies for cancer in general and melanoma specifically should therefore use pre-clinical assays that incorporate the physiological complexity of TME and capture the interplay between stroma cells in the TME and cancer cells. Drug testing using engineered three-dimensional (3D) tissue models that mimic the TME has emerged as a promising human-based assay platform for cancer drug discovery and development [6,7]. Three-dimensional tissue constructs are generated by culturing human cells under conditions that recapitulate the architecture and microenvironment of native tissues or organs, including the complex intercellular interactions, extracellular matrix composition and mechanical properties found in the TME in vivo. A key advantage of these models is their capacity to more accurately predict human responses to candidate compounds, as human-derived cells retain their native genetic makeup and cell-stroma interactions that regulate responses to therapeutics [8,9].

For drug-screening applications, engineered 3D tissue models built in multiwell plate formats enable high-throughput screening, offering the robustness and scalability needed to test compounds across a range of concentrations and assess their potency and efficacy. When biofabricated in a 96-well plate format, these models can support the screening of approximately 1,000 compounds. Here, we describe the development and application of a 3D-bioprinted, full-thickness melanoma skin model for a multi-tiered high-throughput drug screen. Drawing on advances in tissue engineering and fluorescence-based imaging, we developed an assay that measures melanoma tumor growth within full skin tissue models. These models consist of a dermis populated with primary dermal fibroblasts and an epidermis containing primary neonatal keratinocytes, with tumor-like melanoma clusters positioned at the dermal-epidermal junction. Melanoma cells and dermal fibroblasts were transduced to express red and green fluorescent proteins, respectively, and confocal microscopy was used to measure tumor growth alongside fibroblast viability as a counter screen. Using this system, we identified compounds that selectively reduce melanoma cell viability while sparing the surrounding stromal cells. Together, this approach provides a versatile assay platform that incorporates the TME for melanoma therapeutic discovery, with the potential to accelerate the identification of clinically relevant treatments.

## Materials and Methods

### Cell Culture

Neonatal human dermal fibroblasts (HDF; Zen-Bio, Cat# DFN-F) and A375 melanoma cells (ATCC, Cat# CRL-1619) were cultured at 37 °C and 5% CO_2_ in Dulbecco’s Modified Eagle Medium (DMEM; Gibco, Cat# 11965) supplemented with 10% fetal bovine serum (Cytiva, Cat# SH30071.03) and 1% penicillin-streptomycin (Gibco, Cat# 15140). Prior to expansion, HDF and A375 cells were fluorescently labeled by transduction with zsGreen and tdTomato lentiviral particles, respectively (Takara, Cat# 0038VCT and 0036VCT), at a multiplicity of infection of 20 with a 4-hour incubation. The labeled cells were then expanded in culture before cryopreservation.

Neonatal normal human epidermal keratinocytes (NHEK; ScienCell, Cat# 2100) were cultured at 37 °C and 5% CO_2_ in keratinocyte medium (Lonza, Cat# 192060). HDF, A375, and NHEK cells were cryopreserved in aliquots of 4 × 10^6^ cells at passages 5, 9, and 3, respectively. For downstream applications, aliquots were thawed into culture flasks 2 days prior to use, and cells were harvested at 60-80% confluency.

### Dermal Hydrogel Preparation

Gelatin (Sigma-Aldrich: Cat# G1890; final concentration 4.5% w/v) was dissolved in a fibrinogen solution (20 mg/mL stock in 0.87% NaCl; Sigma-Aldrich: Cat# F3879; final concentration 7.7 mg/mL) at 37 °C. Following complete dissolution, collagen type I (Corning: Cat# 354249; final concentration 4 mg/mL) was added and mixed thoroughly. The mixture was supplemented with 10× phosphate-buffered saline (PBS) to balance ionic strength and brought to final volume with sterile water. The hydrogel was maintained at 37 °C prior to use and neutralized with 1 N NaOH immediately before addition of HDF cells.

### Biofabrication of Three-Dimensional Melanoma Skin Constructs

HDF were detached using 0.05% trypsin (Gibco: Cat# 25300), pelleted by centrifugation, and resuspended in the dermal hydrogel preparation at a concentration of 2 × 10^6^ cells/mL. The cell-hydrogel mixture was loaded into a glass syringe (RegenHU: Cat# 900013152) and chilled on ice 1-2 minutes to accelerate polymerization. A 21-gauge needle (Nordson: Cat# 7018233) was attached, and the syringe was mounted on a RegenHU 3DDiscovery bioprinter (RegenHU, Switzerland).

The dermal hydrogel with HDF cells was bioprinted on day minus-7 on top of a membrane (8.0 μm pore polyester) insert of a 96-well transwell plate (Corning: Cat# 3384) as a double layer covering the transwell membrane, with edge wells excluded to minimize evaporation artifacts. Each layer of the construct was printed as previously described, approximately 0.3mm thick in a circular outline tracing the edge of the membrane, with serpentine filling perpendicular between the two layers. [10]

Immediately following printing, 250 µL of DMEM (Gibco: Cat# 11965) containing thrombin (0.5U/mL; Sigma: Cat# T6884) was added to the basal compartment of the receiver plate to crosslink the fibrinogen component of the hydrogel. After a 1-hour incubation at RT, medium was replaced with complete DMEM (10% FBS, 1% penicillin-streptomycin, 0.025 mg/mL aprotinin: Sigma-Aldrich: Cat# A4529), and constructs were maintained at 37 °C with media changes every 2 days.

Epidermalization media and cornification media were formulated as previously described. [11]. On day 0, medium was changed to 350uL epidermalization medium. NHEK and A375 cells were harvested using 0.05% trypsin, combined at a 50:1 ratio (NHEK: A375), pelleted, and resuspended in keratinocyte medium. The mixed cell suspension was pipetted onto the bioprinted dermal constructs at a density of 19,000 cells/well. Constructs were incubated for 4 days with basal media changes every 2 days. On day 4 the tissues were raised to the air-liquid interface (ALI) using a custom 3D printed adaptor made of SBS-format compliant material (Figure S1). At this time, the medium formulation was changed to cornification medium, and the volume was adjusted to 350 µL per well to maintain the tissues at the ALI. Cornification medium was changed daily at the same volume for the remainder of the assay.

### Drug Screen Protocol

350 nL of compound solutions in DMSO at 10 mM were spotted into receiver base plates (Corning: Cat# 3382) with an Echo acoustic dispenser and stored at -80 °C until use. Compound well mapping was blinded for all three rounds of high-throughput screening (HTS). Positive controls for each round were 10 µM bortezomib in 3 wells per plate, and negative control consisted of three wells with 0.1% DMSO included in each plate. Before use, compound plates were thawed at room temperature for at least 30 min. Compounds were resuspended in 350 µL/well of cornification medium prior to assembling with tissue inserts. Drug-containing medium was changed daily for 6 days, and constructs were fixed on day 14. For dose response testing, dose responses were first prepared in DMSO and then spotted as described above.

### Fluorescence Imaging and Data Acquisition

Constructs were imaged live on day 8 (prior to drug treatment) and following fixation on day 14 (after 6 days of compound exposure). For the primary and secondary screening rounds, imaging was performed using a Molecular Devices ImageXpress. Fluorescent images were acquired with a 4× objective at excitation/emission wavelengths of 475/650 nm (zsGreen) and 544/570 nm (tdTomato), and image analysis performed using PerkinElmer Columbus software. For the tertiary screen, imaging was performed on a Revvity Opera Phenix system using a 5x objective with factory settings for FITC and tdTomato channels, with image analysis on Revvity SimA software. Z-stacks were collected at 25 µm step intervals transversing the tissues, with image analysis performed on the maximum projection.

### Data Processing and Normalization

Total red channel fluorescence intensity per well was used as an indicator of melanoma cell growth, while total green channel fluorescence intensity per well served as an indicator of fibroblast viability. To normalize well-to-well differences in number of tumors, melanoma growth was quantified as the ratio of fluorescence intensity on day 14 relative to day 8, normalizing each well’s response to its own tumor number baseline at the onset of drug treatment. Drug response was calculated using the following equation:

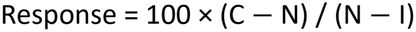

where C = C_14_/C_8_ (day 14/day 8 ratio for each test well), N = N_14_/N_8_ (ratio for the untreated negative control), and I = I_14_/I_8_ (ratio for the bortezomib-treated positive control). Each well yielded two independent response values, one for each fluorescence channel.

### Clinical Trial Data Analysis

Classification of compounds by melanoma clinical trial history was performed using Anthropic’ s Super Intelligence (SI) assistant, Claude, which reviewed publicly available clinical trial records and literature for each compound through January 2025.

Compounds were designated as “clear success” if they achieved regulatory approval or met primary endpoints in a Phase 3 trial; “partial success” if they demonstrated meaningful but insufficient activity in early-phase trials; “tested but insufficient efficacy” if they failed to meet primary endpoints; and “untested” if no melanoma-specific trial data were identified. These designations represent SI-assisted interpretations of the available literature. A randomly selected subset of the SI-generated classifications of clinical trial success was manually reviewed against published literature and confirmed to be accurate.

## Results

Figure 1 shows the schematic for the full skin thickness melanoma tissue model. The protocol was based on previous work by our laboratory [12]. Briefly, the dermis layer made up of human dermal fibroblasts in a gelatin/fibrinogen/Collagen Type I hydrogel is bioprinted on top of an 8.0 μm pore polyester membrane of a 96-well transwell insert. After 7 days incubation, a suspension of keratinocytes and melanoma cells are pipetted on top of the dermis layer, incubated for 4 days to form a monolayer and then lifted at ALI at Day 4 to initiate epidermis cornification which continues for 10 days until Day 14. Drugs are added basolaterally via the media on day 8, and replaced daily for 6 days prior to fixation.

**Figure 1.**
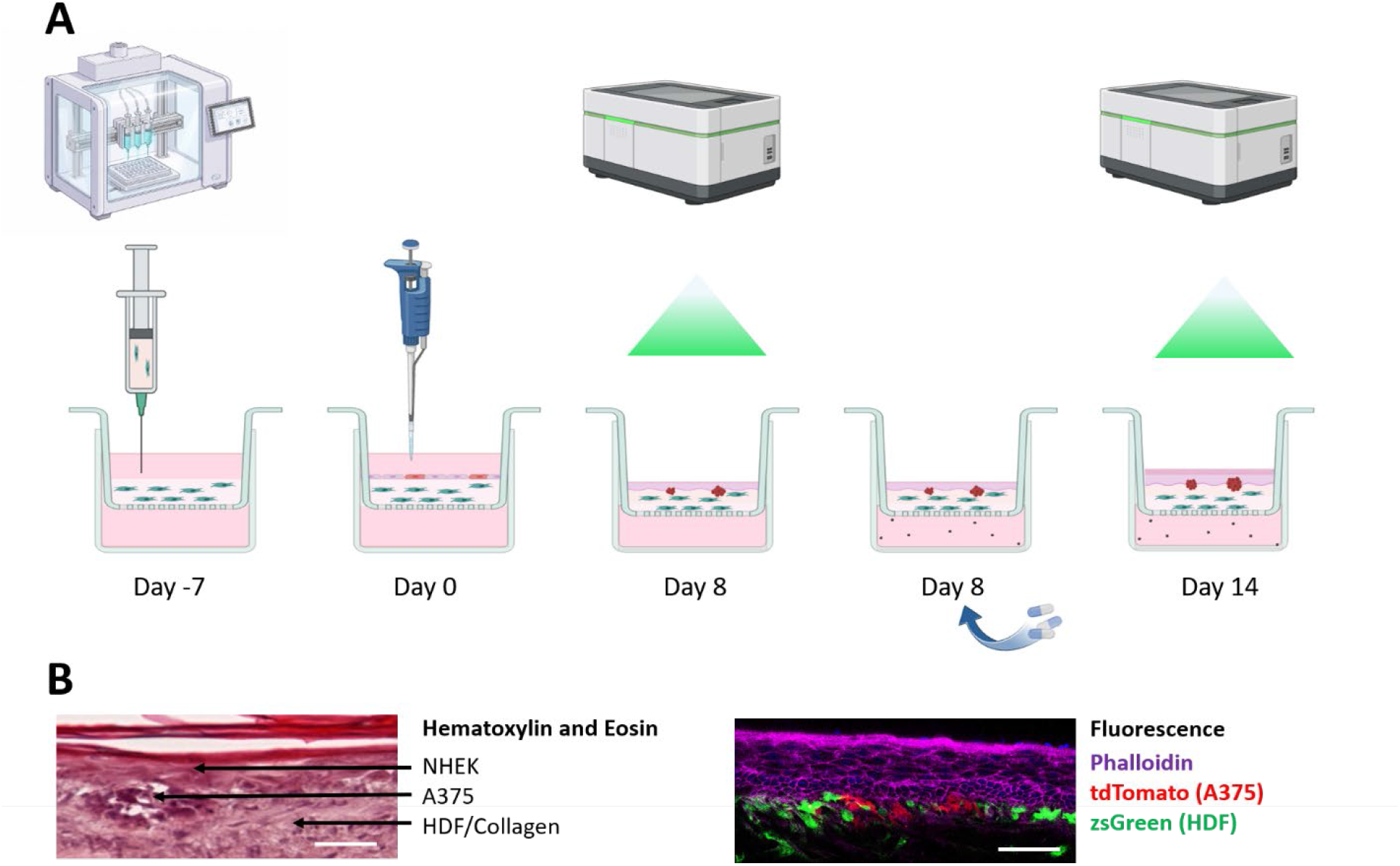
**A)** Schematic of printing protocol. On day -7 hydrogel containing HDF is bioprinted into transwells. On day 0 NHEK and A375 cells are pipetted onto apical side. On day 8 baseline images are acquired with a high content reader, and drugs are added to media on basal side. On day 14 samples are fixed for imaging. Created in BioRender. Derr, P. (2026) https://BioRender.com/lm9ai9p **B)** Histology: H&E and fluorescence imaging. Scale bar 50µm.

To characterize melanoma tumor development within the 3D skin constructs, untreated tissues were imaged at days 1, 3, 6, 8, 10, 12, and 14 (Figure 2). Over the course of the experiment, melanoma cells progressively migrated and coalesced into discrete multicellular clusters, ultimately forming tumor-like aggregates (Figure 2A). Quantitative analysis revealed that the fluorescence intensities of both fibroblasts (zsGreen) and melanoma cells (tdTomato) increased at broadly comparable rates over time, with calculated doubling time of green and red as 3.1 and 2.9 days respectively, at the detectable proliferative state after a lag phase which lasted about 3 days for the A375 cells but only one day for HDF cells (Figure 2B).

**Figure 2.**
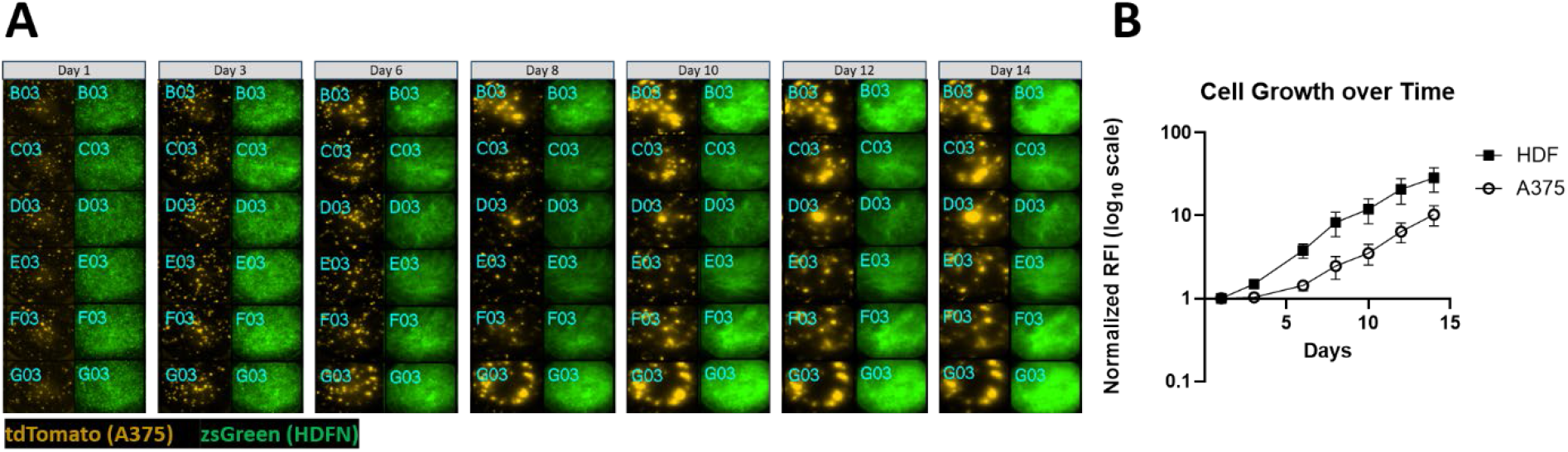
A) Time course images show clustering of A375 cells over time. Image contrast and intensity settings were adjusted independently for each day to optimize visualization and are not directly comparable across images. B) Steady state growth as measured from total fluorescence intensity from the green (HDF) and red (A375) channels and plotted for each time point. Error bars are standard deviation Of N=6 wells per time point. Doubling times were calculated to be around 3 days at the growth phase for both A375 and HDF by fitting to an exponential growth curve.

### Primary screen

A total of 971 compounds from the NCATS Mechanism Interrogation PlatE (MIPE) 5.0 Oncology library were screened at a single concentration of 10 µM as N=1. The selected compounds targeted a broad range of mechanisms of action (MOA) and were mostly in clinical development or clinically approved. Active compounds from the primary screen were selected for secondary testing if they produced greater than 50% reduction in red channel signal (melanoma viability) with less than 20% reduction in green channel signal (fibroblast viability), yielding 177 compounds (Fig 3 and Supplemental Data).

**Figure 3.**
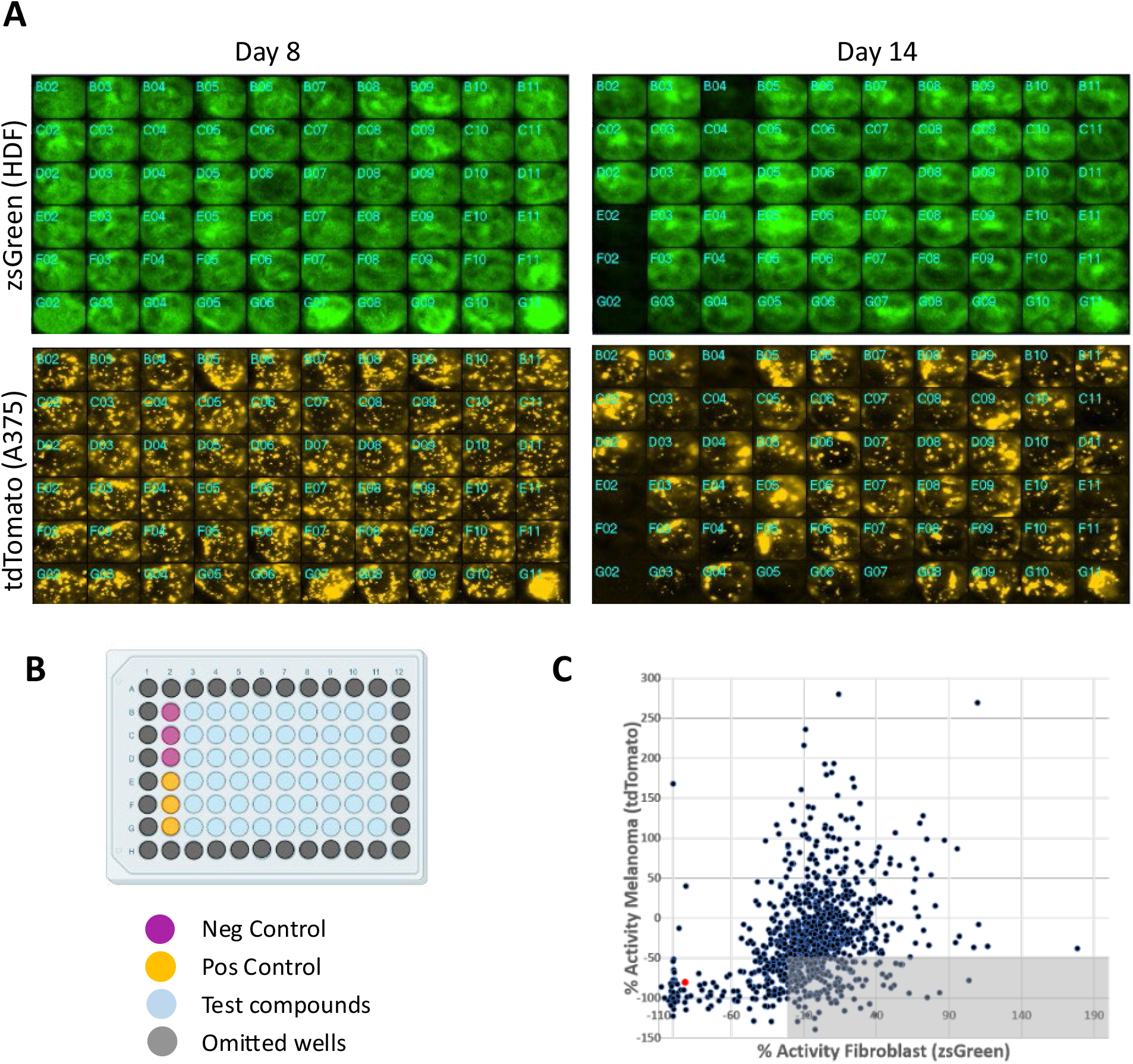
A) Representative images of one plate from primary screen. Omitted edge wells are not shown. B) The plate layout shows location of 3 Negative controls (media + DMSO), 3 positive controls (media +10 µM bortezomib), test compounds (10 µM) and omitted wells containing PBS for humidity control. C) Scatter plot of % activity for the tdTomato (A375 cells cells) vs % activity from the zsGreen (HDF) fluorescence intensity signal. The shaded region indicates compounds meeting hit-selection criteria, defined as >50% reduction in tdTomato signal and <20% reduction in zsGreen signal. The red data point denotes the positive control Bortezomib.

### Drug dose response confirmation testing

Selected compounds were tested in a 6-point dose-response format from 10 µM to 0.04 µM, as N=1 (Figure 4), using the same assay conditions as for the primary screen. Compounds were selected for further testing if they met both of the following criteria: Inhibited % activity for the tdTomato (A375) and zsGreen (HDF) fluorescence intensity signal at the highest concentration (% maximum response) by > 35% and < 20% respectively. Sixty-five compounds satisfied these criteria. To reduce this set to a targeted panel of 27 compounds for additional retesting, final compound selection was based on selecting for broad representation across distinct MOA, as well as a preference for relatively poor performance in a previously conducted internal 2d assay (high AC_50_ values and low % maximum responses). These criteria were designed to enrich compounds demonstrating selective activity in 3D relative to 2D cellular models, thereby highlighting the potential of 3D assays to identify clinically relevant candidates that may be missed in conventional 2D screens.

**Figure 4.**
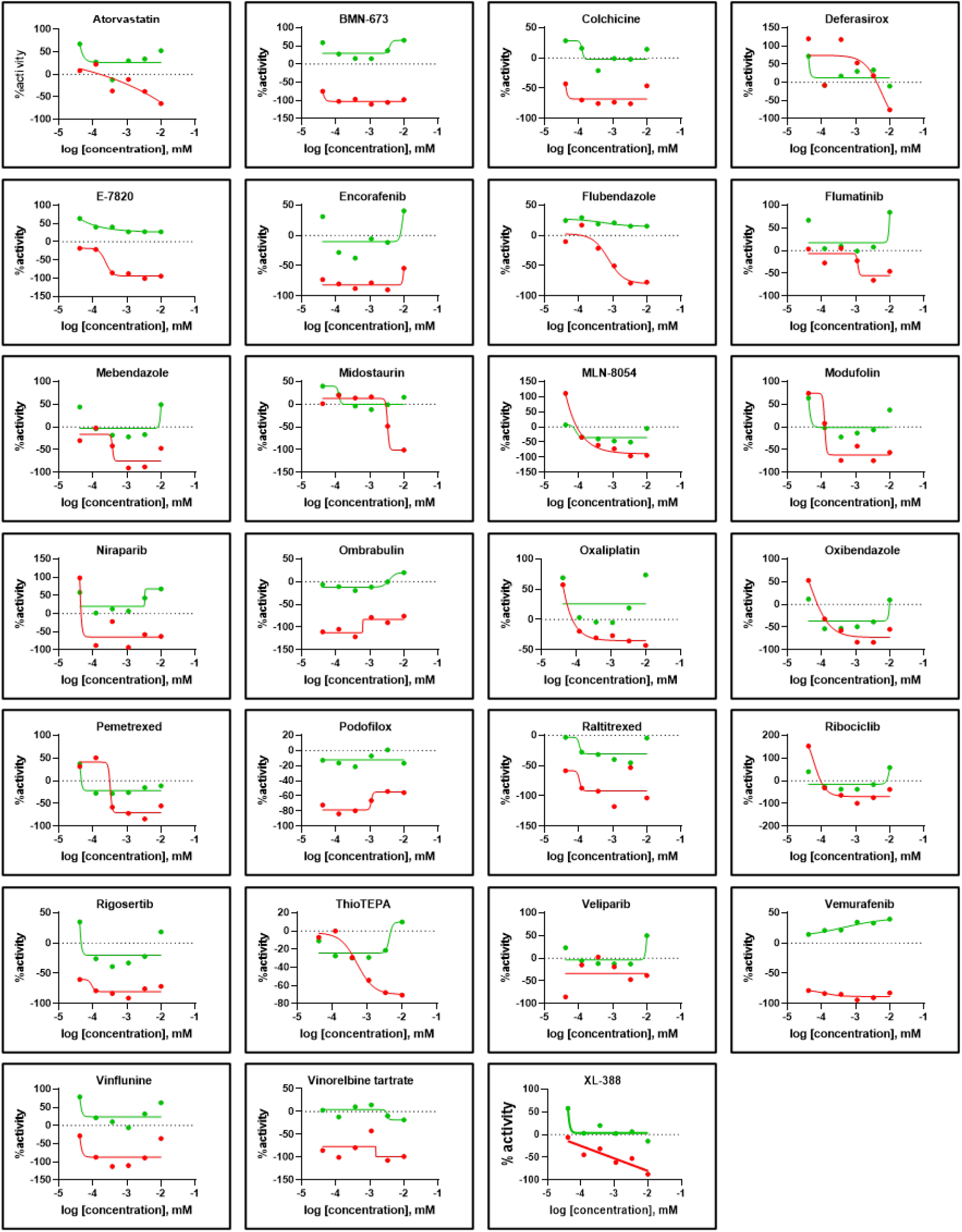
Dose-response curves for selected compounds after first confirmatory test. Red curves represent effects on tdTomato (A375) and green curves on zsGreen (HDF). N=1

Eighteen compounds met the criteria for designation as confirmed hits based on three converging criteria from their dual-reporter dose-response profiles (Figure 5): robust A375 (tdTomato) curve fit quality, substantial maximal cancer cell killing, and a favorable separation between the HDF (zsGreen) and A375 response curves. Across the selected compounds, the green HDF curves were largely flat or modestly declining across the tested concentration range, while the red A375 curves descended steeply, producing a visually consistent and reproducible pattern of cancer-preferential suppression. This separation was quantitatively reflected in the HDF maximum response minus A375 maximum response differential (mean ΔMax = +91.4% for selected vs. +19.4% for non-selected compounds). Selected compounds achieved A375 IC_50_ values in the low sub-nanomolar to low nanomolar range, with particularly well-defined sigmoidal A375 curves observed for E-7820, MLN-8054, Vemurafenib, and Vinorelbine tartrate, where steep red curve descent with concurrent HDF sparing was unambiguous. Several compounds exhibited exceptional selectivity windows: Podofilox, Vemurafenib, Ombrabulin, and Encorafenib displayed among the largest visual and quantitative separations between red and green curves, with HDF-to-A375 IC_50_ ratios ranging from ∼256-to >750-fold. BMN-673 (Talazoparib) and Modufolin similarly showed robust A375 suppression at the lowest concentrations tested, with red curves already suppressed at the leftmost points of the dose range, precluding convergence to a tractable IC_50_ within the assayed window. Rigosertib presented a notably distinct profile in which the HDF curve rose anomalously at higher concentrations while A375 signal remained suppressed, a pattern consistent with a cancer-selective cytotoxic mechanism accompanied by a possible HDF proliferative response. Colchicine displayed a comparable anomalous HDF upturn at high concentrations, while its A375 curve remained robustly suppressed. The nine non-selected compounds were excluded due to one or more of the following: poor A375 curve fit (R^2^ < 0.75), insufficient maximal cancer cell killing, biologically implausible positive A375 bottom values indicative of a net proliferative rather than cytotoxic response, or A375 IC_50_ values outside the screened concentration range. Together, the 18 selected compounds included antimitotics, antifolates, PARP inhibitors, kinase inhibitors, and tubulin-targeting agents, demonstrating potent, well-fitted A375 dose-response curves and relative HDF sparing that collectively justifies their advancement to downstream validation.

**Figure 5.**
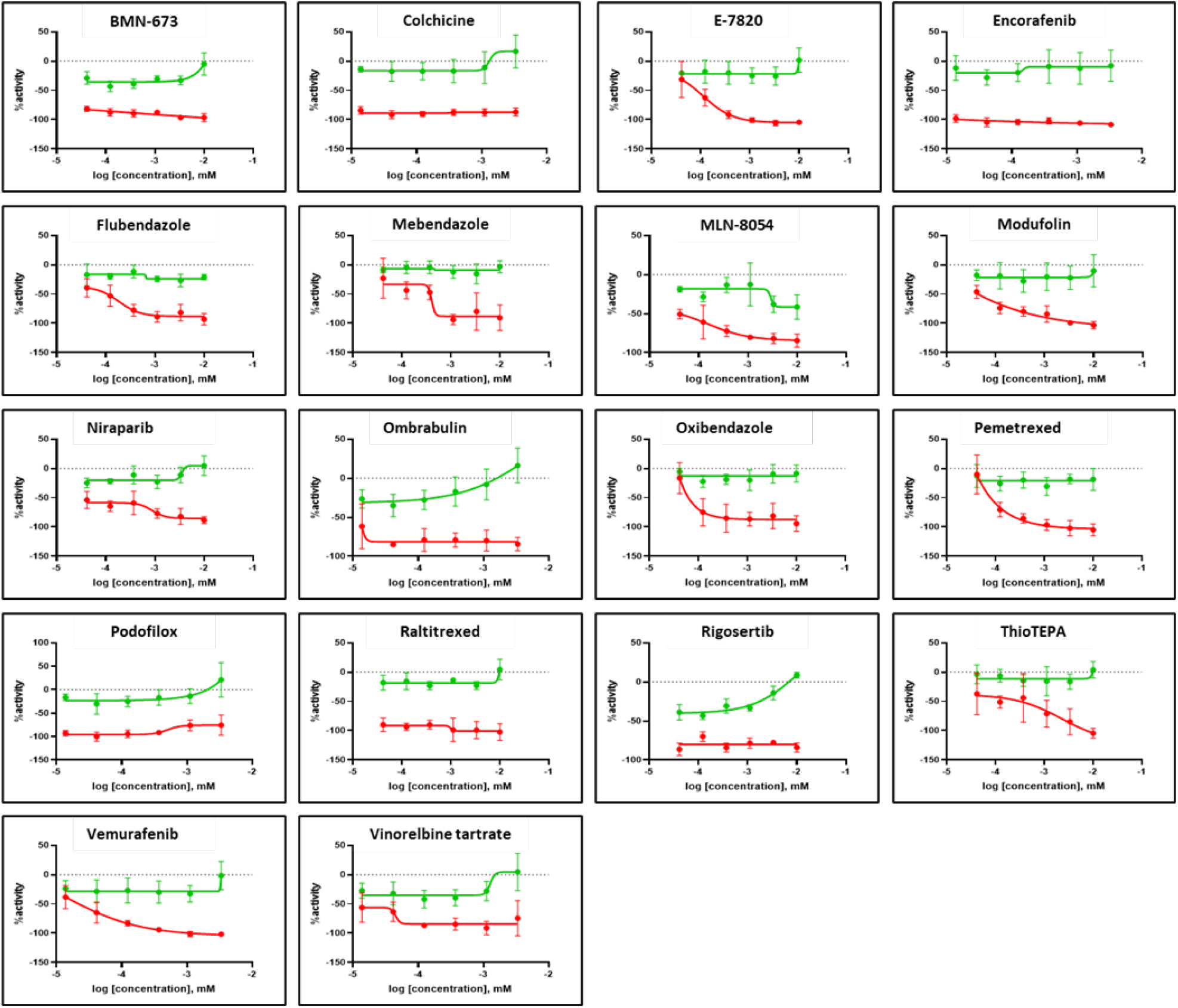
Dose response curves for 18 final selection compounds. Red curves represent effects on tdTomato (A375 cells cells) and green curves on zsGreen (dermal fibroblasts). N=3, error bars indicate standard deviation.

**Figure 6.**
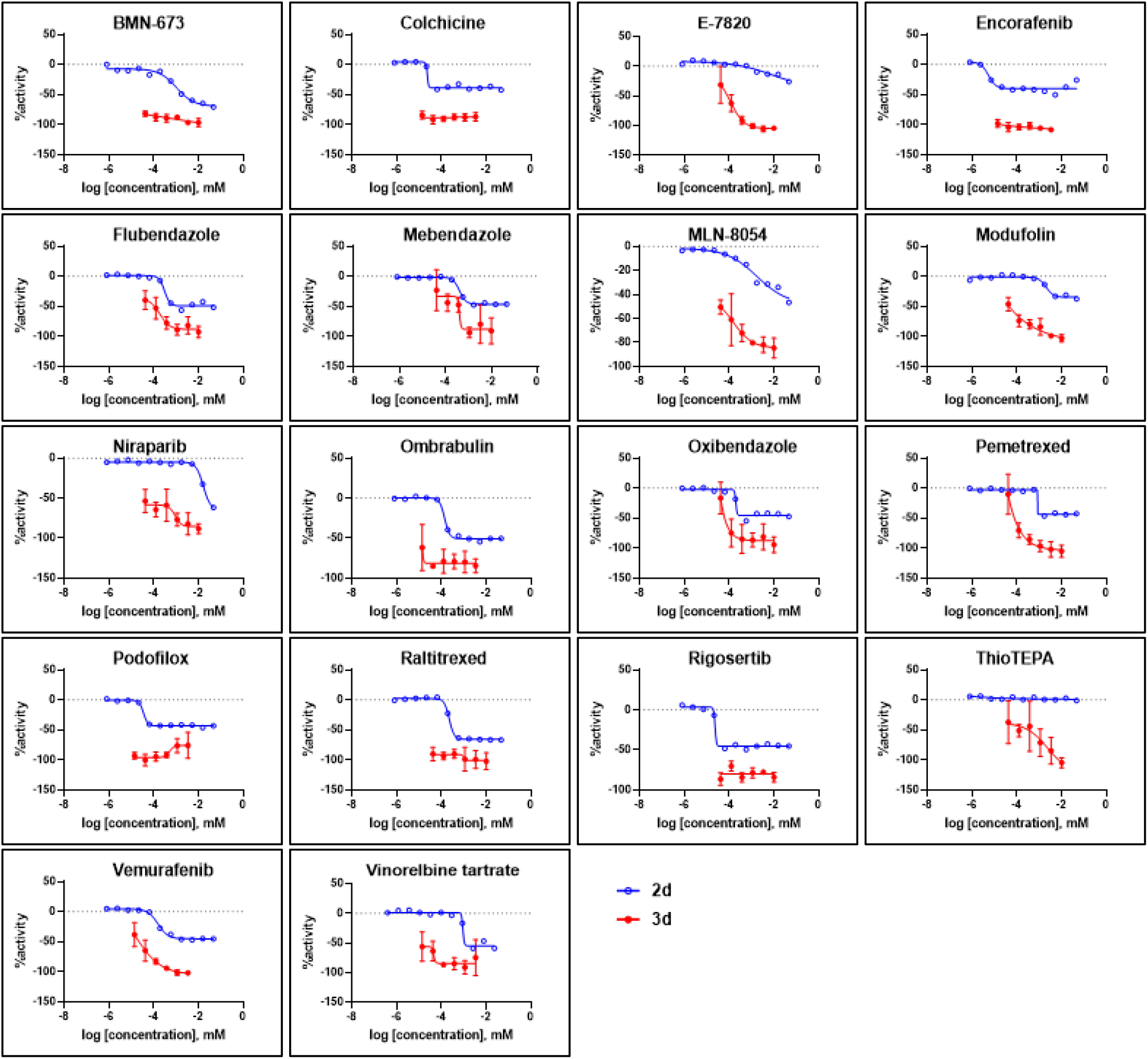
Dose responses of selected compounds for effects on A375 cell growth as measured in 2D assay (2 days, CellTiterGlo) and 3D skin tissue model (6 days, fluorescence intensity). N=3, error bars indicate standard deviation.

### Comparison of 3D and 2D Dose-Response Profiles

Dose-response profiles obtained from the tertiary 3D screen were compared with those from a A375 2D cell proliferation MIPE collection screen conducted previously at NCATS using a 48 hrs., CellTiterGlo cell viability assay readout (see Supplemental Data). Across all 27 compounds evaluated in both formats, the 3D assay system consistently produced higher maximal responses than the 2D assay, and most compounds also exhibited lower IC_50_ values in the 3D melanoma skin model. Furthermore, many of the dose responses for these compounds in 2D show how efficacy plateau at approximately 50% activity, suggesting that in this assay format, most of the compounds appear to be cytostatic, not cytotoxic. We benchmarked the two A375 assays using bortezomib, a proteasome inhibitor included as a cytotoxic positive control. As anticipated, bortezomib produced near-complete cytotoxicity (maximal response approximately 100%) in the A375 2D assay. The systematic elevation of both efficacy and potency in the 3D model relative to the 2D format suggests that the bioprinted melanoma tissue construct provides a more sensitive and potentially more physiologically relevant platform for the identification of active compounds.

### Clinical relevance

Systematic screening of the initial 971-compound primary library using the 3D skin melanoma assay and the analysis criteria described above demonstrated progressive enrichment for melanoma-relevant drugs across successive screening rounds (Figure 7). In the unfiltered library, 15% of compounds had been previously evaluated in melanoma clinical trials, of which 14% were classified as having achieved clear clinical success. This proportion increased substantially with each round of selection: 32% of the 177-compound secondary set had been tested in melanoma trials; 49% in the 71-compound subset; and 56% in both the 27- and 18-compound tertiary sets. Importantly, the success rate among previously tested compounds also improved with successive filtering, culminating in a final 18-compound panel in which 30% of compounds with melanoma trial history achieved clear clinical success and 60% demonstrated combined success (clear success or partial activity). This progressive enrichment validates the multi-tiered selection strategy and confirms effective curation toward compounds with both a higher likelihood of prior melanoma evaluation and a greater probability of demonstrating clinical activity. Notably, the BRAF inhibitors vemurafenib and encorafenib, cornerstone agents for the treatment of BRAF V600, mutant melanoma, were retained through all selection rounds and were present in the final 18-compound panel, providing independent validation of the screening platform.

**Figure 7.**
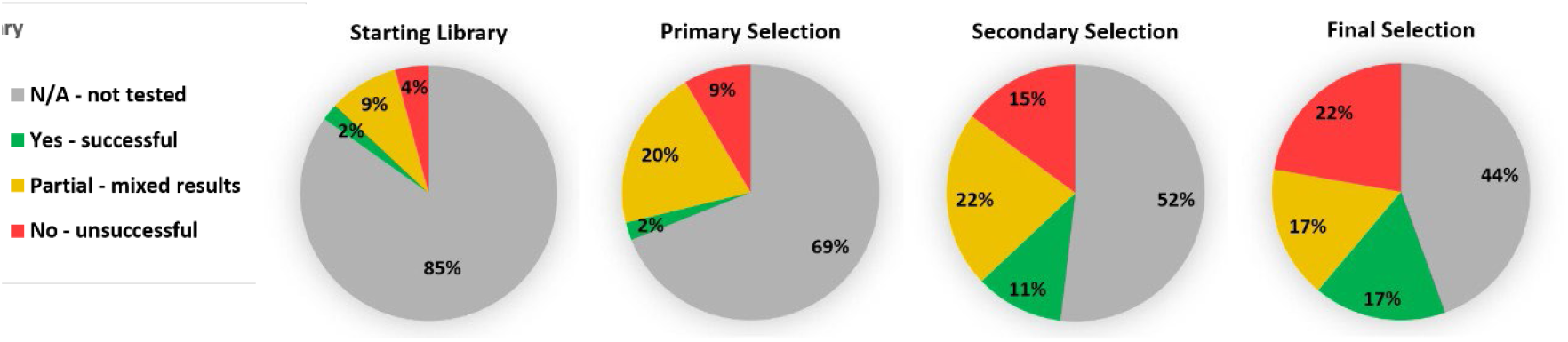
Pie charts depicting the melanoma clinical trial outcomes of selected compounds from each screening phase. Percentages indicate compounds in each round of testing with successful, unsuccessful, partially successful, or no reported melanoma clinical trial outcomes. Data generated using Claude (Anthropic).

Clinical trial data analysis and visualization were performed using Claude (Anthropic), which assessed each compound’s melanoma trial history through January 2025 and generated the comparative figures shown in Figure 7.

The final 18 compounds selected represent a highly curated and mechanistically diverse collection enriched for compounds with demonstrated or potential clinical activity. Three compounds were classified as clear clinical successes: vemurafenib and encorafenib (both BRAF V600 inhibitors, FDA-approved for BRAF-mutant melanoma) and rigosertib (ON-01910), a multi-kinase inhibitor with documented clinical activity for melanoma. Six compounds were classified as partial successes for melanoma, including mebendazole (an anthelmintic agent with emerging antineoplastic properties), thiotepa (an alkylating agent used in high-dose regimens), and vinorelbine (a vinca alkaloid microtubule inhibitor), among others targeting DNA synthesis and repair pathways. Four compounds, pemetrexed (an antifolate) and three kinase/signaling inhibitors (MLN-8054, ombrabulin, and E-7820), were previously tested for melanoma but demonstrated insufficient efficacy as monotherapy in clinic. The eight remaining compounds lack prior melanoma clinical evaluation and include flubendazole and oxibendazole (benzimidazoles of the same class as the partially successful mebendazole) and colchicine (a microtubule inhibitor with established anti-inflammatory activity). The combined success rate of 60% among compounds with prior melanoma testing (10/18 showing any activity) substantially exceeds the baseline rate observed in the unfiltered 971-compound library, validating the progressive enrichment approach and identifying this panel as a high-priority set for further mechanistic and translational investigation.

## Discussion

This study demonstrates the utility of a bioprinted, full-thickness three-dimensional melanoma skin model as a platform for high-throughput drug screening and presents a curated panel of 18 candidate compounds with selective activity against melanoma cells in a physiologically relevant tissue context. Taken together, the findings support three principal conclusions: that 3D tissue models offer meaningful advantages over conventional 2D assays for oncology drug screening; that the progressive multi-tiered selection strategy employed here effectively enriches for clinically relevant compounds; and that the identified candidates, including both approved therapeutics and repurposing opportunities, warrant further mechanistic and translational investigation.

A central finding of this work is the systematic divergence between compound responses in the 3D and 2D assay formats. Across all 18 compounds evaluated in both systems, the 3D model consistently yielded higher maximal responses and, in most cases, lower IC_50_ values than the corresponding 2D assay. This divergence is consistent with recent work in other tumor types: using organotypic microtumor slices from breast and pancreatic cancers, Nishida-Aoki et al. reported that roughly three times as many compounds were active in 3D tissue than in matched 2D cultures and identified stroma-directed hits that were entirely missed by conventional cancer-cell screens. [13] The convergence of this finding across independent platforms, tumor types, and model formats reinforces the conclusion that 2D screens systematically underestimate the activity of compounds whose efficacy depends on the tissue context. The deliberate selection of compounds showing higher potency and efficacy in the 3D system relative to 2D, one of the explicit criteria used to advance compounds from secondary to tertiary screening, underscores this point. By prioritizing 3D-selective activity, the screening strategy enriched for compounds capable of overcoming the microenvironmental barriers that contribute to treatment failure in vivo and may have de-prioritized compounds whose apparent activity in 2D does not translate to more complex tissue settings. This design principle has implications beyond the present study: it suggests that incorporating 3D models earlier in drug discovery pipelines could reduce late-stage attrition by filtering out compounds with activity that is unlikely to be translated to the clinic.

Further mechanistic studies will be needed to study whether these differences in 2D vs 3D pharmacology reflect fundamental differences in the tumor microenvironment between formats. Conventional monolayer cultures lack the structural complexity, cell-cell contacts, extracellular matrix interactions, and diffusion gradients that characterize solid tumors in vivo, and these features are known to influence drug penetration, metabolism, and cellular stress responses [14]. Three-dimensional tumor models, by contrast, more faithfully recapitulate the spatial organization, cell-cell and cell-ECM interactions, and the microenvironmental heterogeneity of native tumors that are central to drug resistance mechanisms [15].

The progressive enrichment of melanoma-relevant compounds across successive screening rounds provides strong evidence for the validity and discriminating power of the platform. The proportion of compounds with prior melanoma clinical trial evaluation increased from 15% in the unfiltered library to 56% in the final 18-compound panel, while the combined success rate among tested compounds rose from 14% to 60%. Most compellingly, the BRAF inhibitors vemurafenib and encorafenib, two of the most clinically important agents in the current standard of care for melanoma, were independently retained through all selection rounds and identified as top hits. The rediscovery of established, FDA-approved melanoma therapies from a library of nearly 1,000 diverse compounds, without any prior knowledge embedded in the selection algorithm, constitutes a meaningful internal validation of the screening approach.

Among the most intriguing findings is the emergence of benzimidazole anthelmintics as a compound class of interest. Mebendazole, a broad-spectrum antiparasitic agent, was identified as a partial success in the screen and is mechanistically related to flubendazole and oxibendazole, two additional benzimidazoles present in the untested portion of the final panel. Benzimidazoles exert antineoplastic effects in preclinical models, in part through disruption of tubulin polymerization, a mechanism shared with colchicine and the established chemotherapeutic agent vinorelbine, both of which also appeared in the final panel [16][17]. The convergent identification of multiple tubulin-targeting agents through an unbiased screen suggests that this mechanism of action may be particularly relevant in the 3D melanoma tissue context, possibly reflecting the importance of cytoskeletal dynamics in tumor invasion and cluster formation observed in this model. Mebendazole has attracted increasing preclinical and clinical interest as a drug-repurposing candidate across multiple cancer types. Its favorable safety profile, well-characterized pharmacokinetics, and low cost make it an attractive candidate for further evaluation in melanoma. [18]. The co-identification of rigosertib, a multi-kinase inhibitor with documented clinical activity, further broadens the mechanistic diversity of the candidate panel and highlights the complementary value of kinase pathway inhibition alongside microtubule-targeting strategies.

A few limitations of the present study warrant consideration. First, all experiments were conducted using a single melanoma cell line (A375), which harbors the BRAF V600E mutation and represents only one of several molecularly distinct melanoma subtypes. While the use of A375 cells enabled direct comparison with prior internal 2D screening data and provided a well-characterized baseline, it limits the generalizability of the findings to BRAF wild-type, NRAS-mutant, NF1-mutant, and triple-wild-type melanoma subtypes, which collectively account for a substantial proportion of clinical cases and exhibit distinct drug sensitivity profiles [19, 20]. Future studies should incorporate additional cell lines representative of melanoma’s molecular diversity, as well as patient-derived melanoma cells, to assess the breadth of the platform’s predictive utility.

Second, the model as constructed does not incorporate immune cell populations, which are now recognized as central determinants of melanoma behavior and treatment response. The dramatic clinical success of immune checkpoint inhibitors in melanoma has established beyond doubt that tumor-immune interactions profoundly shape disease progression and therapeutic outcomes [21]. The absence of T cells, natural killer cells, macrophages, and other immune effectors from the current construct means that the platform is not suited to evaluate immunotherapeutic agents or to model the immunosuppressive tumor microenvironment that drives resistance to both targeted therapy and immunotherapy. This represents a significant gap relative to the current clinical landscape of melanoma treatment. The biofabrication protocols developed, though, will enable the incorporation of additional relevant cells, and our group has already published protocols for the incorporation of macrophages in full thickness skin tissue models [22].

Third, while the enrichment of clinically validated compounds provides indirect evidence for the platform’s translational relevance, none of the newly identified candidate compounds have yet been validated in animal models or clinical settings. The relationship between activity in the 3D assay and in vivo efficacy remains to be established, and it is possible that some hits will not translate despite their selectivity in the tissue model. Rigorous in vivo validation of the top candidates is an essential next step before any clinical inference can be drawn.

Finally, the comparable apparent growth rates of the two cell populations warrant cautious interpretation. Although melanoma cells would be expected to proliferate more rapidly than fibroblasts embedded within a 3D hydrogel matrix, the two populations exhibited similar doubling times (2.9 and 3.1 days, respectively). One possible explanation involves the 8 µm pore size of the transwell membrane: fibroblasts positioned at the base of the hydrogel construct may migrate through these pores and colonize the underside of the insert, gaining access to a proliferative niche unavailable to the melanoma cells. The observed increase in fibroblast fluorescence may therefore reflect a combination of in-situ growth and pore-mediated migration, rather than in-situ proliferation alone, whereas the increase in melanoma fluorescence likely reflects accelerated tumor growth driven by cluster-induced spatial compaction and the inherently higher proliferative capacity of malignant cells. Importantly, this consideration affects only the baseline growth characterization of the untreated constructs; it does not confound the drug-response measurements, which are calculated as normalized day-14/day-8 fluorescence ratios within each well and are therefore internally controlled for baseline signal.

### Future Directions

The findings of this study suggest several productive directions for future investigation. Most immediately, the 18 confirmed hits, particularly the eight compounds lacking prior melanoma clinical evaluation, represent high-priority candidates for mechanistic characterization and in vivo validation. Compounds such as flubendazole, oxibendazole, and colchicine, which share mechanistic class with active hits but lack melanoma-specific clinical data, are especially compelling repurposing candidates given their established safety profiles in other indications.

With respect to model development, the incorporation of immune cell populations represents the most significant opportunity to extend the platform’s biological relevance. Integration of autologous or allogeneic T cells, macrophages, or dendritic cells into the tissue construct would enable evaluation of immunotherapeutic agents and permit investigation of the complex reciprocal interactions between tumor cells, stromal fibroblasts, and immune effectors that govern melanoma progression. Similarly, the addition of vascular elements, either through endothelial cell incorporation or microfluidic perfusion systems, would more accurately recapitulate the tumor vasculature and enable the study of drug delivery dynamics, including the penetration barriers that are central to solid tumor pharmacology.

Beyond melanoma, the modular architecture of the bioprinting platform described here is, in principle, adaptable to other epithelial malignancies that arise within stratified tissue contexts. Application of analogous models to cutaneous squamous cell carcinoma, Merkel cell carcinoma, or other skin-associated tumors could leverage the existing fabrication and screening infrastructure while broadening the oncological scope of the approach. More broadly, the demonstration that a tissue-engineered 3D model can prospectively enrich for clinically active compounds supports the wider adoption of such platforms in oncology drug discovery, with the potential to improve the efficiency of the development pipeline and reduce the high rates of late-stage clinical trial failure that continue to constrain the field [23, 24].

In summary, this work establishes a bioprinted three-dimensional melanoma skin model as a viable and discriminating platform for high-throughput oncology drug screening, identifies a curated panel of candidate compounds with selective antimelanoma activity, and highlights the value of integrating 3D tissue models into early-stage drug discovery. The progressive validation afforded by multi-tiered screening, combined with the independent rediscovery of approved melanoma therapies, supports the translational potential of this approach and provides a foundation for the continued development of more complex, immunocompetent tissue models for cancer drug discovery.

## Acknowledgements

This work was funded by the NIH Intramural Research Program (IRP) and the Cure Acceleration Network program at NCATS. The contributions of the NIH author(s) are made as part of their official duties as NIH federal employees are in compliance with agency policy requirements and are considered Works of the United States Government. However, the findings and conclusions presented in this paper are those of the author(s) and do not necessarily reflect the views of the NIH or the U.S. Department of Health and Human Services.

## CRediT Author Statement

Conceptualization: PD, KD-M and MF; Methodology: KD-M, SF, TV, PD; Investigation: KD-M, PD; Formal analysis: MS, PD, KD-M, TV and MF; Visualizations: PD, KD-M and MF; Writing - Original Draft: PD; Writing - Review & Editing: PD, KD-M, SF and MF; and Supervision, Project Administration, and Funding: MF

