## Supplementary material for "A Three-Dimensional Bioprinted Skin Melanoma Model for High-Throughput Drug Screening": Figure S1

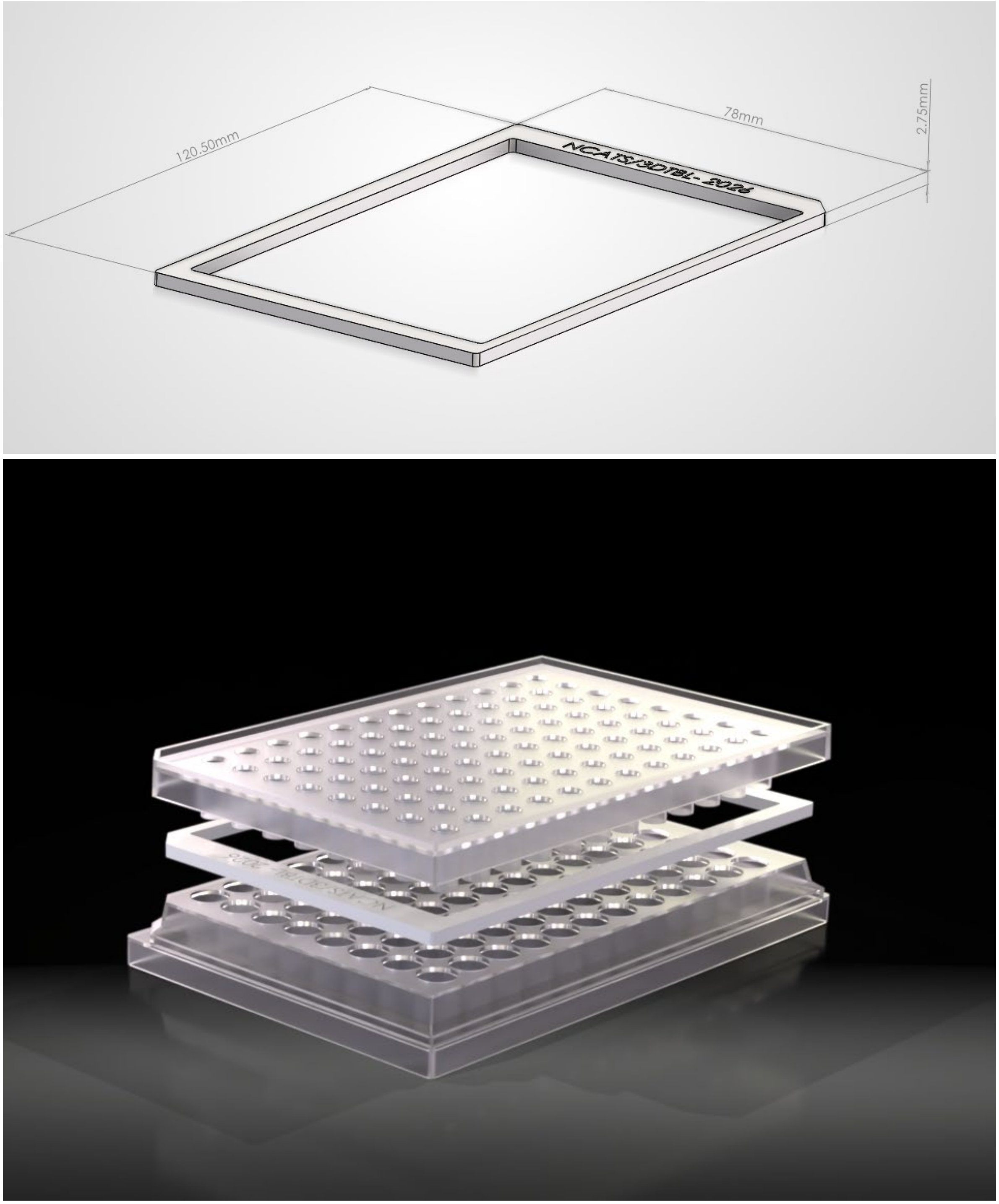


**Figure S1:** Dimensions and rendered assembly of 3D printed transwell spacer to allow for increased media volume. Spacer is in SBS-compliant format and made of biocompatible material.
